# ‘A Generalized Reinforcement Learning-Based Deep Neural Network (GRL-DNN) Agent Model for Diverse Cognitive Constructs

**DOI:** 10.1101/2022.06.17.496500

**Authors:** Sandeep S. Nair, Vignayanandam R. Muddapu, C Vigneswaran, Pragathi P. Balasubramani, Dhakshin S. Ramanathan, Jyoti Mishra, V. Srinivasa Chakravarthy

## Abstract

Human cognition is characterized by a wide range of capabilities including goal-oriented selective attention, distractor suppression, decision making, response inhibition, and working memory. Much research has focused on studying these individual components of cognition in isolation, whereas in several translational applications for cognitive impairment, multiple cognitive functions are altered in a given individual. Hence it is important to study multiple cognitive abilities in the same subject or, in computational terms, model them using a single model. To this end, we propose a unified, reinforcement learning-based agent model comprising of systems for representation, memory, value computation and exploration. We successfully modelled the aforementioned cognitive tasks and show how individual performance can be mapped to model meta-parameters. This model has the potential to serve as a proxy for cognitively impaired conditions, and can be used as a clinical testbench on which therapeutic interventions can be simulated first before delivering to human subjects.

## 1. INTRODUCTION

High-level human cognition consists of a variety of functions or capabilities, including selective processing of goal-relevant information, suppression of goal-irrelevant information, action selection, reward processing, working memory, etc. There is a long history of empirical research that studies the various cognitive functions individually while excluding other functions. However, to understand these cognitive functions as functions of an integrative agent, it is essential to study them holistically, revealing the synergies among these functions that come into play as an agent interacts with its environment.

While empirical research on cognitive functions suffers from this fragmented approach due to several challenges including participant burden, limited resources and expertise, theoretical investigation also often reflects this piecemeal approach, offering a wide variety of models that describe individual cognitive functions. There have been efforts to construct integrative computational frameworks that capture a range of cognitive functions. For example, the “ACT-R” system is a general framework for modeling a wide variety of cognitive processes (Anderson, 1997). Subsequently, it was extended to include visual attention, and its properties like speed and selectivity as they vary from subject to subject (Anderson, 1997). Similarly, the “Soar” architecture can successfully integrate different levels of reasoning, planning, reactive execution, and learning from experience (Laird, 2018). The importance of more holistic and integrative models has been emphasized by several researchers, resulting in many unified computational models of cognition. As these models evolved, a certain similarity among these modeling architectures began to reveal itself. For example, common features of three such cognitive architectures viz., ACT-R (Anderson, 1997), Soar (Laird, 2018; Young & Lewis, 2012), and SIGMA (Rosenbloom, Demski, & Ustun, 2016) have been described (Laird, Lebiere, & Rosenbloom, 2017).

Neuropsychiatric disorders are characterized by a wide range of cognitive dysfunctions, and the degree to which these disorders are mapped to specific neural substrates is still being resolved (Millan et al., 2012). Just as theoreticians sought to create unified architectures of cognition, experimental cognitive scientists also made efforts to move away from exclusivist approaches and began to study multiple cognitive functions in human population cohorts simultaneously (Weintraub et al., 2013). One such experimental system is the *BrainE* platform, which includes a range of cognitive assessments like selective attention(SA), response inhibition(RI), working memory(WM), and distractor processing(DP) in both non-emotional and emotional context (Balasubramani et al., 2021). The *BrainE* system measures both behavioral parameters and electroencephalography (EEG) signals, thereby creating an opportunity to relate cognitive behavior to neural substrates.

In this paper, we present a unified architecture of cognition that can model a range of cognitive functions, including SA, RI, WM, DP, etc. The proposed model has the following components: 1) sensory representation, 2) memory, 3) value computation, 4) exploration, and 5) action selection. The model is cast broadly within the framework of reinforcement learning (RL) (Chakravarthy, Joseph, & Bapi, 2010; Chakravarthy & Moustafa, 2018; Shivkumar, Muralidharan, & Chakravarthy, 2017; Sridharan, Prashanth, & Chakravarthy, 2006). Notably, the action selection strategy, which involves pursuit of explorative and exploitative modes, each of which is regulated based on the underlying value dynamics is novel to our approach. The model has elements common to deep neural networks and two novel neural elements that are not typically found in such networks viz., 1) flip-flop neurons and 2) oscillator neurons. First proposed in (Holla & Chakravarthy 2016), the flip-flop neurons are fashioned after flip-flops in digital systems theory and can store memories. In the oscillator network, the lateral interactions are designed such that the oscillators exhibit desynchronized dynamics. Such oscillator networks have been used before to implement exploratory functions essential for achieving randomness in action selection in RL models (Chakravarthy and Balasubramani 2014). We hypothesized that this modeling framework can replicate the subject’s performance with respect to diverse cognitive decision-making tasks.

In the following section, we will describe the methods starting with a brief overview of the experimental setup, the cognitive testing paradigms used, and the various tasks conducted to evaluate cognitive abilities. This is followed by a brief overview of the model architecture that is used to simulate the experimental tests, various building blocks of the model architecture and their mathematical formulation, and how they are integrated to mimic an experimental subject. In the results section, we summarize the main results, including the training phase, the performance evaluation and the meta-parameters used for tuning the performance. We compare the model performance with experimental results. The final section concludes the results and discusses the utility, limitations, and future scope.

## 2. MATERIALS AND METHODS

The proposed Generalized Reinforcement Learning-based Deep Neural Network (GRLDNN) agent model, as shown in the Figure 1, can simulate various experimental paradigms that can test different cognitive functions such as SA, RI, WM, and DP.

**Figure 1.**
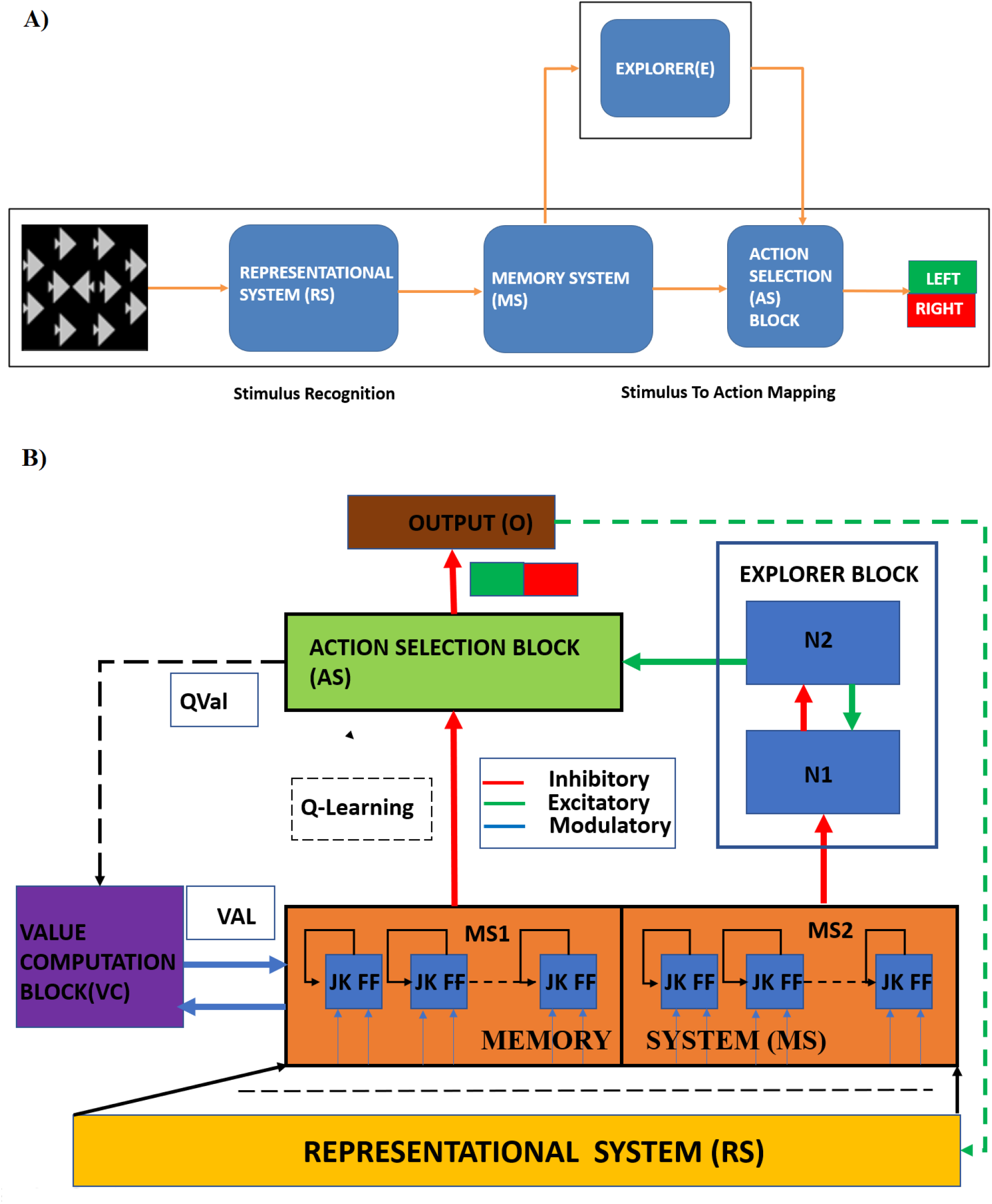
**A) Overview of (Generalized Reinforcement Learning-based Deep Neural Network) GRLDNN model architecture**. RS, Representational System is used for stimulus recognition; Memory System (MS) and Action Selection (AS) block along with the Explorer (E) is used for stimulus to action mapping. The encoder output from RS is presented to the MS where the stimulus is processed, and the action selection takes place at AS. B**) Block diagram of the Agent Model architecture**. RS, Representation System Block; Explorer Block consisting of N1/N2 Oscillator pair; AS, Action Selection Block; O, Output block; VAL, Value Computation Block where Value function is computed; QVal, Q-value function; MS, Memory System block

### 2.1. The Experimental Tasks

A repertoire of tasks was prescribed to capture a range of cognitive functions that comprise human decision-making. The cognitive functions that we focus on in these tasks are the ability to selectively attend to relevant stimuli, inhibit responses to irrelevant stimuli, avoid distractors during an attention task, and working memory. The details of the tasks are described in supplementary section S1.

### 2.2. GRLDNN (Generalized reinforcement Learning-based deep neural network) Agent Model

We present a unified RL-based deep network architecture that can simulate all the experimental tasks described in the previous section.

A schematic of the model architecture is given in Figure 1. The model has five distinctly identifiable components viz. - 1) Representational System (RS) consists of a series of layers – convolutional layers followed by fully-connected layers, that generate compact representations of the input images. 2) Memory System (MS) is a layer of flip-flop neurons that receives the inputs from the RS via a fully connected weight stage. This system has the memory property. 3) Value Computation System (VC) combines the neural outputs of the MS and computes the value function. 4) Explorer comprises a nonlinear oscillator network that interacts via inhibitory connections, generating desynchronized oscillations. The randomness inherent in the chaotic oscillatory dynamics of this system introduces a level of randomness in the action selection at the output layer. 5) Action Selection System (AS) is the output of the entire architecture that combines the outputs of the MS via a trainable weight stage and the output of the Explorer. The AS is trained using Q-learning described in detail in the supplementary materials (De Oliveira, Bazzan, Da Silva, & Grunitzki, 2018).

#### 2.2.1 Representational System

The input stimulus is presented as an image to the input layer of the RS, which is trained as a convolutional autoencoder (Lindsay, 2021; Nerurkar, Chandane, & Bhirud, 2019). The RS module consists of an input layer followed by four convolutional and max-pooling layers. A fully connected layer follows the four layers of the convolutional and max-pooling layers. Another fully connected later is used to reduce the encoder output to 1×64. The convolutional layers use a 3×3 filter window size. Mean squared error is used as the output loss. At the decoder end, the 1×64 feature output is expanded, followed by deconvolutional layers and pooling layers, at the end of which the original image is reconstructed back. Output from the fully connected layer of the encoder part of RS is provided as the input to the MS.

#### 2.2.2 Memory System

The output of the RS module is presented to the MS via a fully connected weight stage. The MS, as mentioned before, is a 1D layer of flip-flop neurons. This layer is divided into two equal sections - MS1 and MS2. MS1 has D1-type flip-flop neurons, whereas MS2 has D2-type flip-flop neurons. The flip flop architecture is provided in the supplementary material (S1.1.1).

The feature vector from the RS module (*X*_*RS*_) reaches the flip-flop neurons of MS1 and MS2 over the weight stages 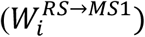 and 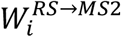, respectively. So, the effective input received at MS1 is 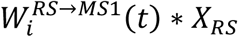 (Figure S4 in the supplementary material).

The output of the encoder part of the RS module is presented as the input, *X*_*RS*_(*t*), to the J and K ports of the flip-flop neurons present in the MS1/MS2 sub-blocks of the MS, 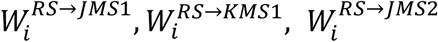 and 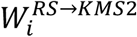 denote the weights from the RS layer to the respective J and K inputs of the MS1/MS2 sub-blocks (Figure 2B).

**Figure 2:**
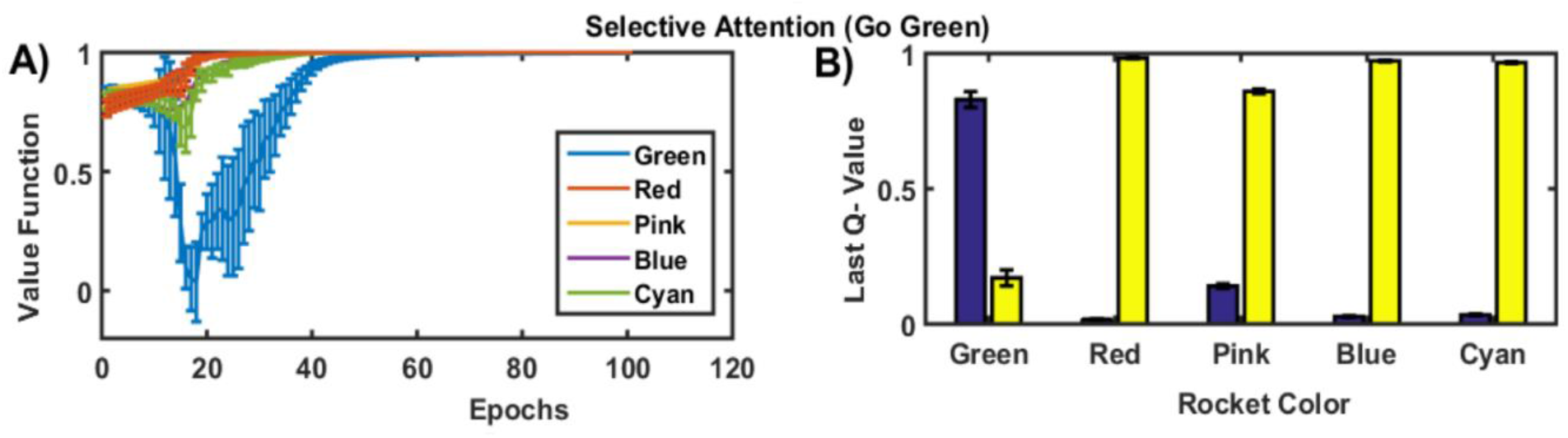
A) The state value functions over the epochs during the training phase of the Go Green (SA) task. B) The Q-value at the end of training for the Go Green task that requires selective attention to the green colored rockets while ignoring other isoluminant colors - red, pink, blue, cyan. The blue bar represents the ‘Go’ action and the yello’ bar represents the ‘No-Go’ action.

##### Computations in the MS1/MS2 Sub-Blocks of the Memory System

The J and K inputs for the flip-flop neurons of MS1 and MS2 are given by equations (1-4) below. The output of the flip-flop neuron is expressed by equations (5-6), which is in line with the circuit diagram and the truth table given in the supplementary section (S1.1.1). The output is also influenced by the modulatory input received from the VC (Sutton & Barto, 2018).

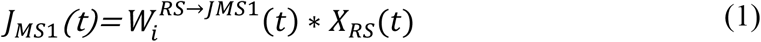

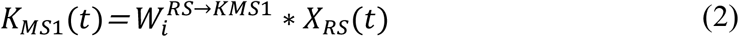

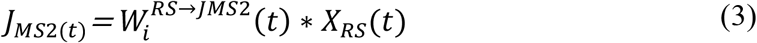

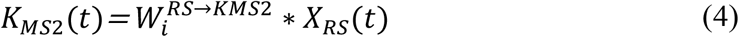

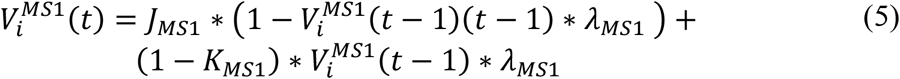

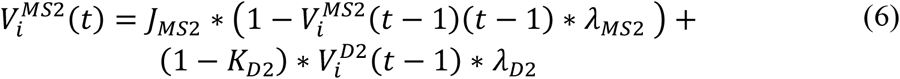

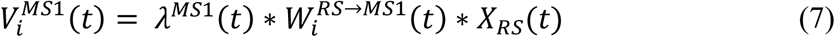

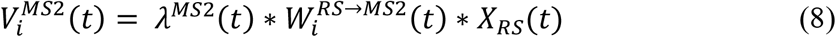

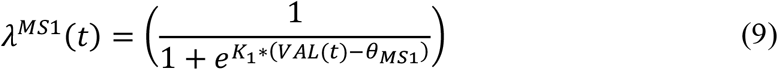

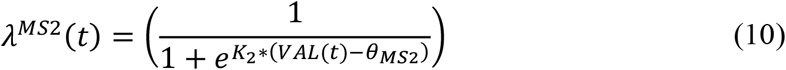

where *K*_1_ < 0 and *K*_2_ > 0 and *λ*^*MS*1^ *and λ*^*MS*2^ are the sigmoid gain parameters.

#### 2.2.3 Value Computation System

The value is computed using a weighted sum of the outputs of the flip-flop neurons of MS1. Thus, the value function ‘VAL’, is computed as per Equation (11) below,

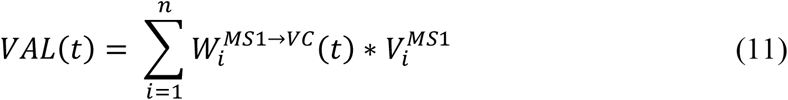

#### 2.2.4 Explorer Module

The explorer module consists of a network of nonlinear oscillators. These are thought to be implemented by two pools of neurons, N1 and N2, connected back-to-back. The MS2-type flip flop neurons of MS project to N1, whereas the output of the N1 neural layer, in turn, influences the N2 neural layer. N1 and N2 form a loop, with inhibitory projections from N1 to N2 and excitatory projections from N2 to N1. Such excitatory-inhibitory pairs of neurons pools have been shown to exhibit oscillations (Chakravarthy & Moustafa, 2018b; Gillies, Willshaw, Atherton, & Arbuthnott, 2002). In the present case, it turns out that the equations that couple a single N1 neuron bidirectionally to a single N2 neuron can be classified as a general oscillator system called Lienard system, which exhibits limit cycle oscillations (Kawahara, 1980). The dynamics of N1-N2 neuronal pools is defined as,

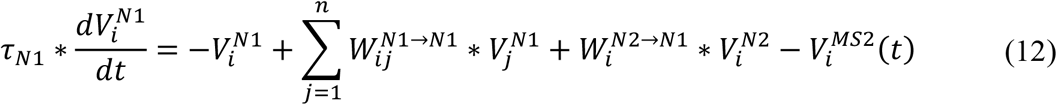

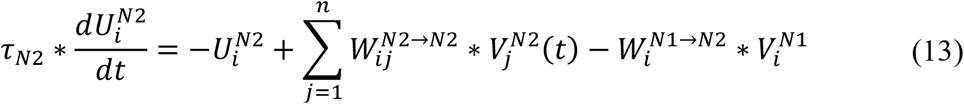

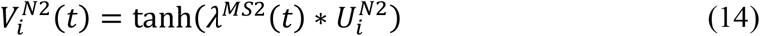

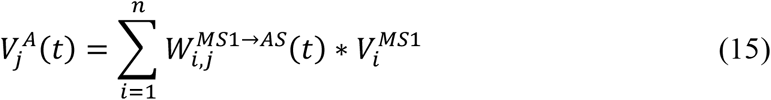

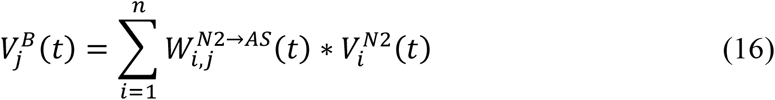

where 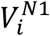 and 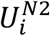 are the internal states of N1 and N2 neurons, respectively, 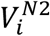 is the output of the N2 neuron, *W*^*N*1→*N*1^ and *W*^*N*2→*N*2^ are weight kernels representing lateral connectivity in N1 and N2 modules, respectively, *τ*_*N*1_ and *τ*_*N*2_ are the time constants of N1 and N2, respectively, *W*^*N*1→*N*2^ is the connection strength from N1 to N2, *W*^*N*2→*N*1^is the connection strength from N2 to N1, and *λ*_*N*2_ is the parameter which controls the slope of the sigmoid in 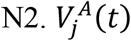 *and* 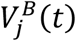 are the inputs arriving at AS1 block from MS1 and MS2 (via N1 &N2) respectively.

##### Lateral Connections among Coupled Oscillators of the Explorer Module

The lateral connectivity in the N1 or N2 network is modeled using constant weights where weights between two neurons is given by *W*^*N*1→*N*1^ and *W*^*N*2→*N*2^,

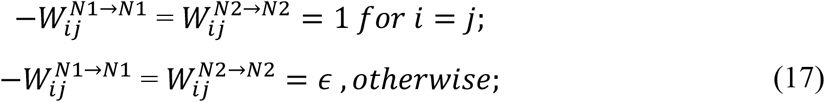

where *ϵ* is the magnitude of the connection strength of lateral connections among the neighbouring neurons in both N1 and N2 modules.

#### 2.2.5 Action Selection Module – Q-learning

The network’s final output is a linear sum of the output of the MS1 block and the output of the explorer module.

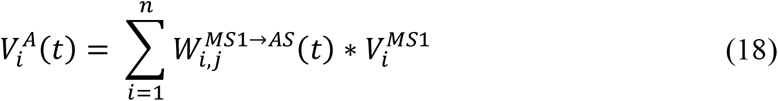

The output 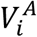 is combined with the output 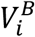 (Equation 16) at the AS block as shown in (Equation 19) below.

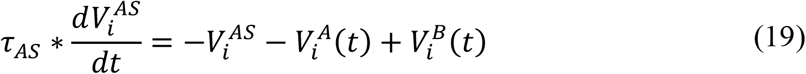

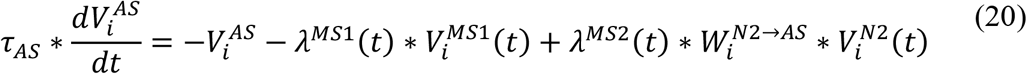

The action selection mechanism at AS is facilitated using a race model (Mandali, Rengaswamy, Chakravarthy, & Moustafa, 2015; Packard & Knowlton, 2002; Y. Smith, Bevan, Shink, & Bolam, 1998; Vickers, 1970). The output of the AS block 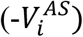 is compared against a threshold *V*_*th*_. The output of whichever neuron ‘i’ crosses the threshold first, it is considered a winner, and the *i*^*th*^ action is selected.

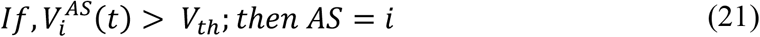

##### Reward and Learning

The weights between the neurons in the MS1 block and the AS module are updated using Q-Learning (Rice & Stocco, 2017) as shown in (eqns. 22 - 24).

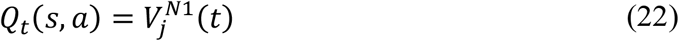

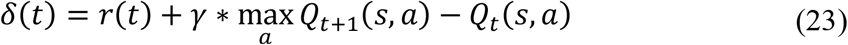

where *γ* = 0 (discount factor).

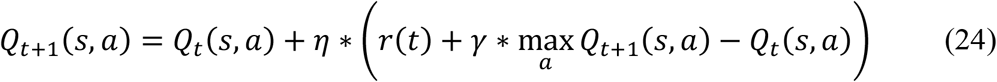

#### 2.2.6 Backward Propagation

In this model, trainable weights are located in three areas: i) MS1-block to AS weight stage, ii) MS1 block to VC block weight stage, and iii) RS to MS1/MS2 block weight stage (Figure S3). The weights between the various modules are updated using the backpropagation algorithm. The weight update between the MS1 and the AS blocks is governed by Q-learning, while the weight update between MS1 to VC is done using temporal difference (TD)-learning. The MS1/MS2 subblocks used flip-flop neurons, and the weight update between RS and MS is done using TD-learning. The weight training equations are described in detail in the supplementary section (S2).

### 2.3. Performance Assessment

The model performance is assessed in terms of the metrics of accuracy, reaction time, speed, consistency, and efficiency, as defined in Supplementary section S3.1.

## 3. RESULTS

In this section, we describe the performance of the GRLDNN model that is used to simulate the four cognitive functions of SA, RI, WM, and DP experimentally assessed using the *BrainE* platform. In the subsequent subsections, we will first show the progress of state value functions and the Q-value functions during the learning process. Then we will show the impact of parameter tuning with respect to lateral connections strengths and threshold (*ϵ* and *V*_*TH*_). Then we implement an inverse model using a neural network that can calibrate the meta parameters of the GRLDNN model so that the model can simulate the experimental performance. We then present a comparison of the average performance between both the model and the experimental results.

### 3.1 Training Phase

As the learning progresses, the magnitude of the value functions at the end of each trial keep increasing and approach the maximum value of 1, as the model learns the task accurately. The Q-values for the respective state and action pairs corresponding to the correct action are higher, whereas the values corresponding to the wrong action are lower. Fig. 2A represents the state value function for the *Go Green* (SA) task, and Fig. 2B represents the Q-value at the end of training epochs, at the last instance of the *Go Green* trials. The state value and Q-value functions for other tasks are described in the supplementary section.

The test dataset contains 33% of the green-colored rockets and 67% of other colored rockets in the SA task. The blue bar represents the ‘Go’ action and the yellow bar represents the ‘No Go’ action.

### 3.2. Effect of Parameter Tuning

The performance of cognitive tasks depends on various factors. We observed variations in the model performance by tuning specific meta parameters, using performance metrics of accuracy, reaction time, speed, consistency, and efficiency. The meta-parameters that are varied are *ϵ* and *V*_*TH*_.

- *ϵ* influences the *lateral connectivity strength* of the N1 and N2 coupled oscillator system, which controls the level of exploration in the model.
- *V*_*TH*_ is the *threshold* that appears in the race model, used in the AS module where the action selection occurs, which controls how fast the decision can be made (reaction time).

#### 3.2.1. Effect of the Threshold (*V*_*TH*_) and Lateral Connectivity Strength (*ϵ*) on the Performance

Fig. 3 shows the variation in performance with respect to speed and consistency for various values of lateral connection strength (*ϵ*) and threshold (*V*_*TH*_) illustrated for Go Green (SA) task. *V*_*TH*_ is varied in the range of 0.3 to 0.5 in steps of 0.1 and *ϵ* takes values of 0.01, 0.03, 0.05 and 0.1. Figs. 3A1 & 3A2 show the variation in decision-making *speed* with respect to changes in *ϵ* and *V*_*TH*_. Figs. 3A3 & 3A4 shows a similar variation in *consistency*. From the results, it can be seen that speed is inversely proportional to both the threshold and lateral connection strength.

**Figure 3:**
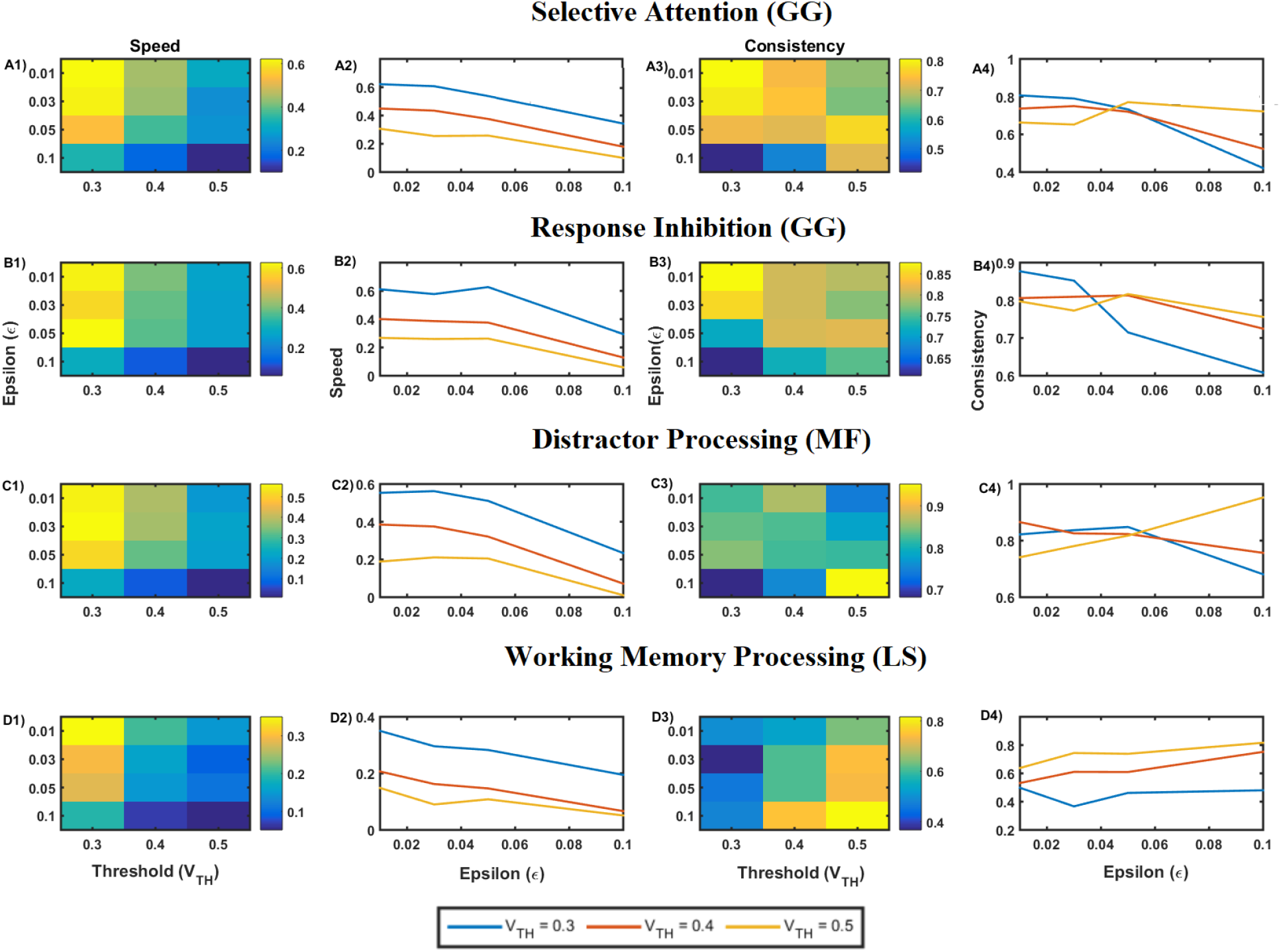
Plot of performance parameters for various tasks, with threshold (V_TH_) varying between 0.3 to 0.5 and lateral connection strength ϵ ranging between 0.01 to 0.1. A1) & A2) The speed with which the decision is made for Go-Green (SA) task, A3) & A4) Consistency for Go-Green (SA) task, which indicates how consistent the performance was with respect to speed across trials. It also indirectly indicates the standard deviation of the speed of performance. Similarly, B1) to B4) represents the Go-Green (Response Inhibition) task, C1) to C4) indicates the Middle Fish (Distractor Processing) task, D1 to D4 indicate performance on the Lost Star task.

Our results show that for lower values of the action selection threshold, the model performance exhibit higher speeds and vice versa. This is expected since the lower the threshold, the faster the action selected. The AS block being implemented as a race model decides on the action to be selected based on the threshold.

Hence by searching for an optimum in the meta-parameter space consisting of lateral connection strength and action selection threshold (*ϵ, V*_*TH*_), it is possible to match the model’s performance with experimental data. We modeled the mapping between the experimental parameters (speed, consistency) and the model meta-parameters (*ϵ, V*_*TH*_), tuned using a simple multilayer perceptron model (MLP). Given the input values of speed and consistency, we can predict the corresponding values of *ϵ* and *V*_*TH*_. Hence by varying the values of the two meta-parameters, we would be able to calibrate the GRLDNN model so that the model can approximately simulate the experimental performance. The model fit between the predicted and desired values of *ϵ* and *V*_*TH*_ is as shown in Fig. 4 below for the *Go Green* (SA) task. Twelve data points were used for training from the combination of meta-parameters (*ϵ* and *V*_*TH*_), and the training error was in the order of ∼10^−5^ after 50000 epochs. The predicted and desired values of both the threshold and epsilon were matched.

**Figure 4:**
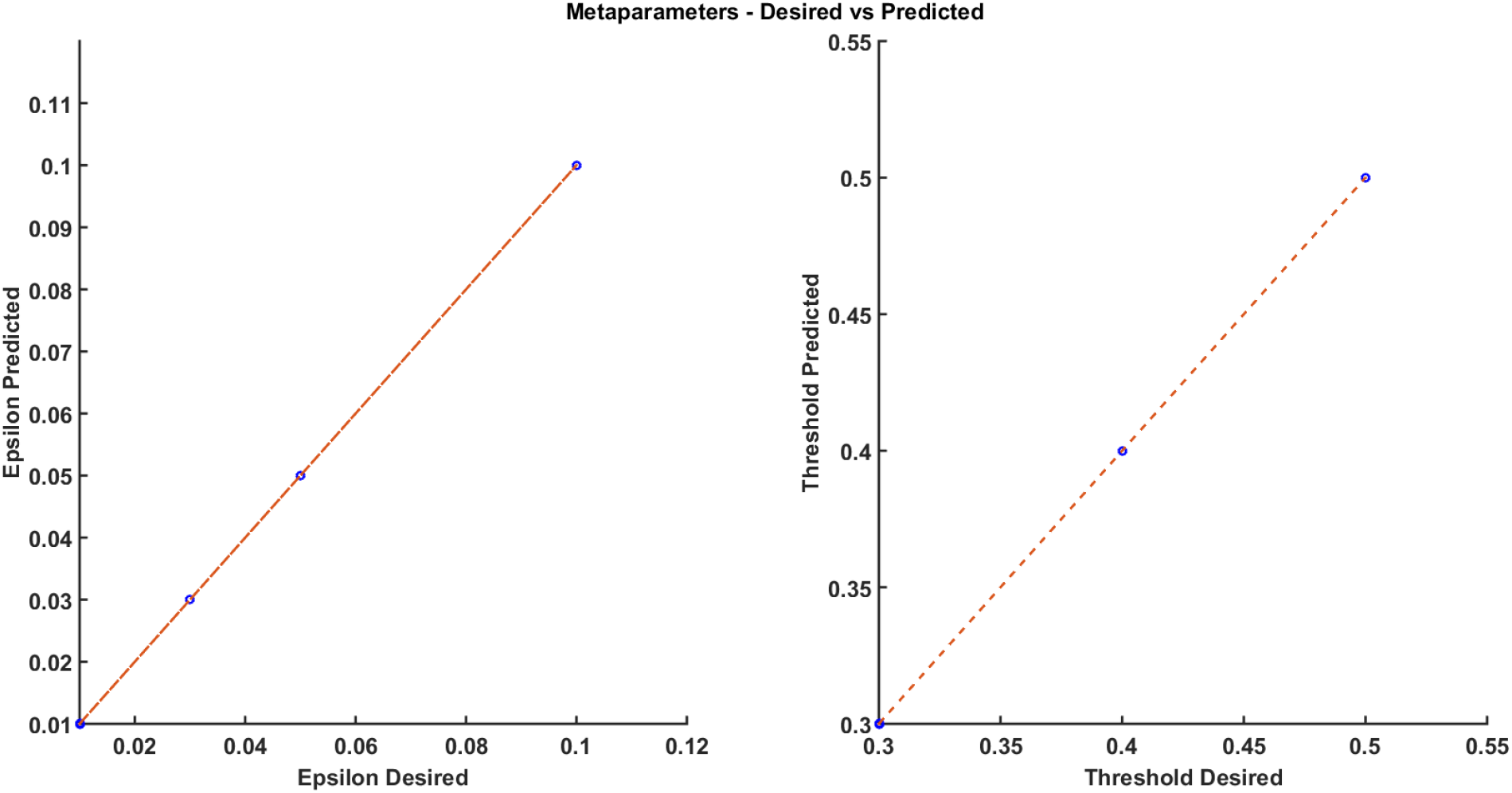
A) The parameter fit is checked for the predicted vs. desired meta parameter (ϵ). B) The parameter fit is checked for the predicted vs. desired meta parameter (V_TH_).

The performance results of the GRLDNN agent model were compared with the experimental results of healthy subjects (Balasubramani et al., 2021). In the GRLDNN agent model, decision-making scenarios were simulated for the different cognitive functions of SA, RI, WM, DP by modeling the test paradigms for *Go Green, Middle Fish*, and *Lost Star* tasks. The model’s performance was tuned using the meta-parameters (*V*_*TH*(*pred*)_ *and ϵ*) to match the experimental results. By navigating through the parameter space as shown in Fig. 3, we selected the values of *V*_*TH*_ = 0.4 *and ϵ* = 0.05, which was found to be closely matching with the average performance results of the experimental subjects. The RMSE (root mean squared error) was found to be lowest at 0.007 for *V*_*TH*_ = 0.4 *and ϵ* = 0.05 when compared with the average performance of experimental results.

Fig. 5 shows the performance comparison of the model and experimental data, where the blue bar represents the model performance, and the yellow bar represents the experimental performance results for all modeled tasks. As seen in the case of the *Go Green* task, the GRLDNN agent model recorded an average speed of 0.3510 ± 0.065 and 0.3756 ± 0.0279 for SA and RI, respectively (Fig. 6A, dark blue bar) compared to the experimental results, which recorded an average speed of 0.3580 ± 0.0534 and 0.3976 ± 0.0612, for SA and RI, respectively (Figure 6A, yellow bar). Similar comparisons can be made for performance metrics for all modeled tasks shown in the figure.

**Figure 5:**
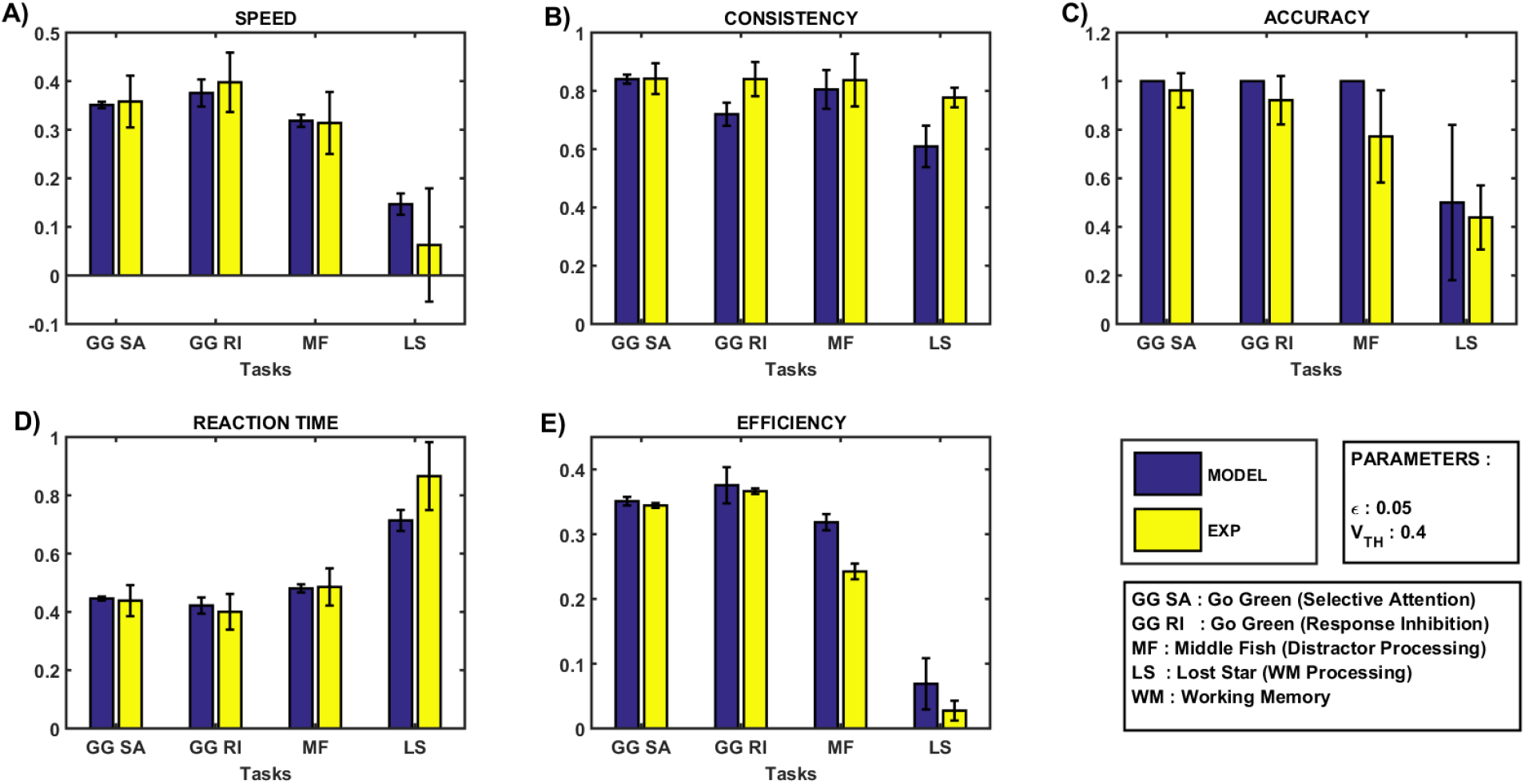
Comparison of performance of the GRLDNN model with the experimental results and data adapted from. (Balasubramani et al., 2021) A) Speed B) Consistency, C) Accuracy, D) Reaction Time, E) Efficiency. EXP, experimental results; MODEL, Model performance Results

**Figure 6:**
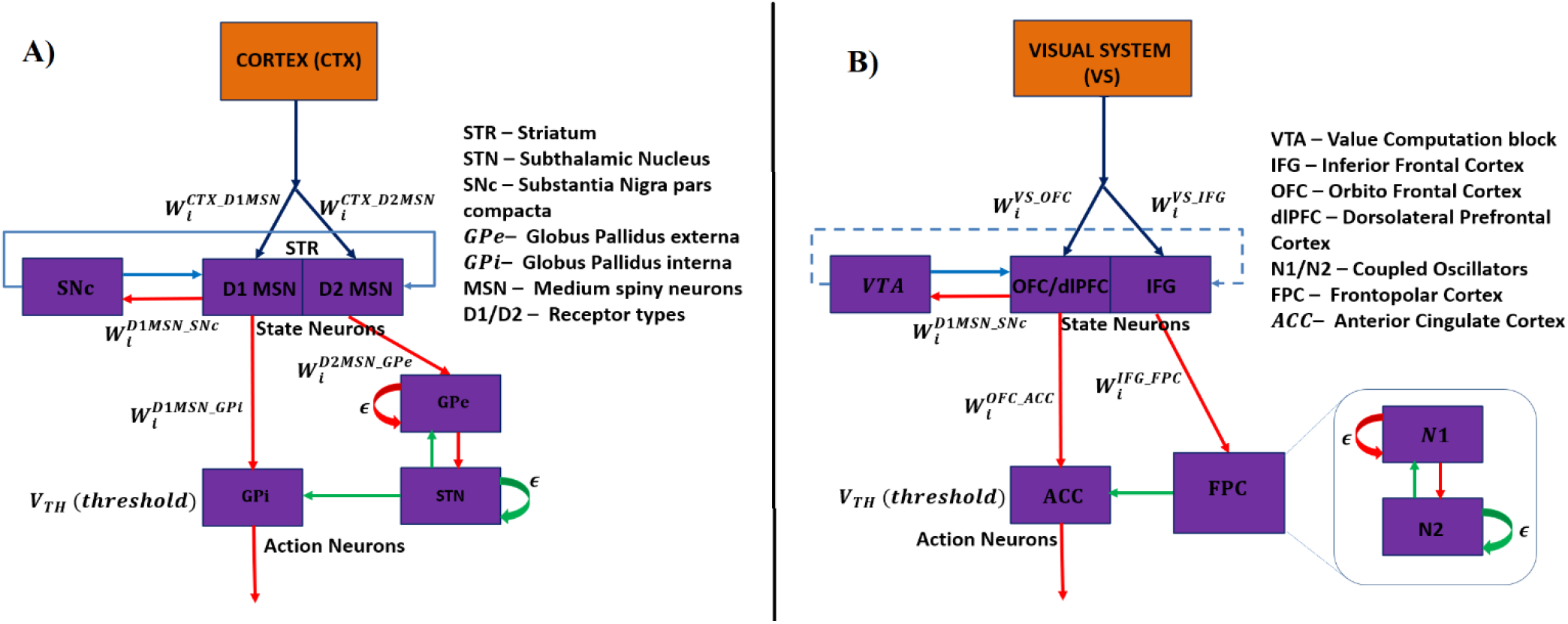
Biological Representation of GRLDNN Agent Model inspired by A) Basal Ganglia (BG) architecture and B) the equivalent representation of prefrontal cortex (PFC) Architecture. SNc, Substantia Nigra pars compacta; STR, Striatum; GPi, Globus Pallidus interna; GPe, Globus Pallidus externa; STN, Subthalamic Nucleus; MSN, Medium Spiny neurons; STN-GPe forms a coupled oscillator system with interconnectivity weights W between STN and GPe and *ϵ* is the lateral connection strengths among the neurons of STN and GPe. Action is selected based on the winner neuron crossing the threshold (*V*_*TH*_) first. VTA, ventral tegmental area; OFC, Orbitofrontal cortex; DLPFC, dorsolateral prefrontal cortex; IFG, inferior frontal gyrus; FPC, frontopolar cortex; ACC, anterior cingulate cortex; FPC is modeled using a coupled oscillator system with interconnectivity weights W between oscillators N1 and N2 and *ϵ* is the lateral connection strengths among the neurons of N2. OFC/DLPFC to ACC weights are updated using Q-learning.

Hence by mapping the performance characteristics across different cognitive tasks onto the meta-parameter space and navigating through the same, we are able to replicate empirical performance on different cognitive abilities using the GRLDNN model.

## 4. DISCUSSION

Using the proposed GRLDNN model, we successfully modeled tasks to assess a variety of cognitive abilities, including SA, RI, WM, and DP. By varying the meta-parameters, *V*_*TH*_ *and ϵ*, we were able to tune the performance outputs of the model (Fig. 3). Notably, our model results were comparable to that of experimental results for healthy subjects (Balasubramani et al., 2021), as shown in Fig. 5.

The current GRLDNN model is an agent model built using a reinforcement learning framework and implemented partly using a deep neural network. Although the model is proposed as a generic agent model that can simulate a variety of cognitive and decision-making tasks, the model’s architecture was originally inspired by an earlier model of the basal ganglia (Chakravarthy and Moustafa 2018) as shown in Fig. 6A. The representational system is analogous to the cortico-striatal projections that are thought to be capable of compressing the cortical state and generate abstract representations of the same (Bar-gad et al 2000). The memory system is analogous to the striatum proper, and the flip-flops are comparable to the medium spiny neurons (MSNs) of the striatum. The MSNs are known to exhibit UP/DOWN states, a property that is thought to subserve working memory functions (Ferbinteanu, 2016; Wilson & Kawaguchi, 1996). In digital systems, flip-flops are used as memory elements that serve as building blocks to implement sequential logic. Thus, in the proposed agent model, the presence of flip-flop neurons in the memory system affords the model the ability to process sequences and perform decision-making functions thereon. The value computation block is analogous to substantia nigra pars compacta (SNc) – it integrates the outputs of the flip-flop neurons of the memory system and computes the value function. The connections from the memory system to the action selection block is analogous to the direct pathway of the basal ganglia. The longer route from the memory system to the explorer block and onward to the action selection block is analogous to the indirect pathway. In modeling literature that describes the decision-making functions of the basal ganglia using reinforcement learning, there is a subclass of models that attribute the role of exploratory drive to the indirect pathway, which is essential to sample the action space randomly (Chakravarthy and Moustafa 2018). Finally, the action selection block itself is analogous to globus pallidus interna (GPi), the output port of the basal ganglia.

The proposed agent model can also be compared to another brain region known for its decision-making functions – the prefrontal cortex (PFC). Fig. 6B shows the analogy between the components of the proposed GRLDNN agent model and areas of PFC whose contributions to decision making have been described extensively (Hare, O’Doherty, Camerer, Schultz, & Rangel, 2008; Klaus & Pennington, 2019; Setogawa et al., 2019; Takahashi et al., 2009).

The dorsolateral prefrontal cortex (DLPFC) receives inputs from the primary and secondary sensory association cortices of the posterior brain (Klaus and Pennington 2019). The DLPFC is also considered to be the terminus of the dorsal visual pathway, also called the “where” or “how” pathway that determines how to use visual information by supplying such information to the decision-making mechanisms of the PFC (Takahashi, Ohki, & Kim, 2013). The DLPFC is also known for its working memory functions, subserved by dopamine-receptor expressing neurons and gated by dopaminergic projections from mesencephalic regions (Klaus and Pennington 2019). Thus, the memory system in the proposed agent model is suitably comparable to DLPFC.

Single unit electrophysiological studies have shown the involvement of the orbitofrontal cortex (OFC) in value computation (Setogawa et al 2019). The role of OFC in value computation was also confirmed by functional imaging studies (Hare et al 2008). Since dopaminergic activity is strongly linked to reward signalling, projections from the ventral tegmental area (VTA) to PFC were implicated in the value computations of OFC (Takahashi et al 2009). Electroencephalographic studies by Bordaud et al (2008), on subjects engaged in decision-making activities, have implicated the frontopolar cortex (FPC) in exploratory behavior. Thus, the exploratory block in the proposed model is comparable to FPC. The inferior frontal gyrus (Aron, Fletcher, Bullmore, Sahakian, & Robbins, 2003; Aron, Robbins, & Poldrack, 2004) is suggested to encode information about NoGo processes and has strong implications for action selection mechanisms, especially action stopping. On the other hand, Anterior Cingulate Cortex (ACC) (Heilbronner & Hayden, 2016; Monosov, 2017) is suggested to encode information about the uncertainty in choices, hence important for estimating utility values of action choices. Currently only one level of working memory processing is tested in the model. Going forward this aspect will be incorporated where the impact of memory load (by increasing the number of stars and the perceptual levels) can be analysed both experimentally as well as in the model.

The current model also can be made more robust and has the scope of conducting patient profiling. By tuning the appropriate model parameters, we are able to match the experimental results, thereby demonstrating its potential use for profiling real patients. There is a scope to explore further and incorporate aspects of various disease conditions and cognitive disabilities into the model. With respect to the modeling of working memory, we have considered tests at only one difficulty level in our model. There is a scope to scale the model to adopt multiple levels of cognitive loads. The future work also includes integrating the cortical and the subcortical modules into a single framework. In the future, these modeling efforts could also be expanded to include emotion processing and modeling of electrophysiological signals. Altogether, this study opens doors to modeling various cognitive dimensions of the same individual through a unified agent-based modeling framework.

## 5. AUTHOR CONTRIBUTIONS

SSN - Conceptualization; Model development; Implementation; Formal analysis; Investigation; Methodology; Validation; Writing – original draft; VRM - Conceptualization; Model development; Implementation; Formal analysis; Investigation; Methodology; Validation; Writing – review & editing; VC - Model development; Implementation; Formal analysis; Investigation; PB - Conceptualization; Formal analysis; Writing – review & editing; Validation; JM - Formal analysis; Writing – review & editing; DR - Formal analysis; Writing – review & editing; VSC - Conceptualization; Model development; Data curation; Formal analysis; Investigation; Methodology; Validation, Writing – review & editing; Supervision.

## Supplementary Material

### S1. The Experimental Tasks

We followed the results from the *BrainE* experimental paradigm for our simulation study and modeled various tasks used in that framework (Balasubramani et al., 2021). Three tasks which were modeled are: *Go Green* (to assess selective attention and response inhibition), *Middle Fish* (to assess distractor processing), and *Lost Star* (to assess working memory) (Figure S1). Cognitive task details are elaborated in the sections below.

**Figure S1:**
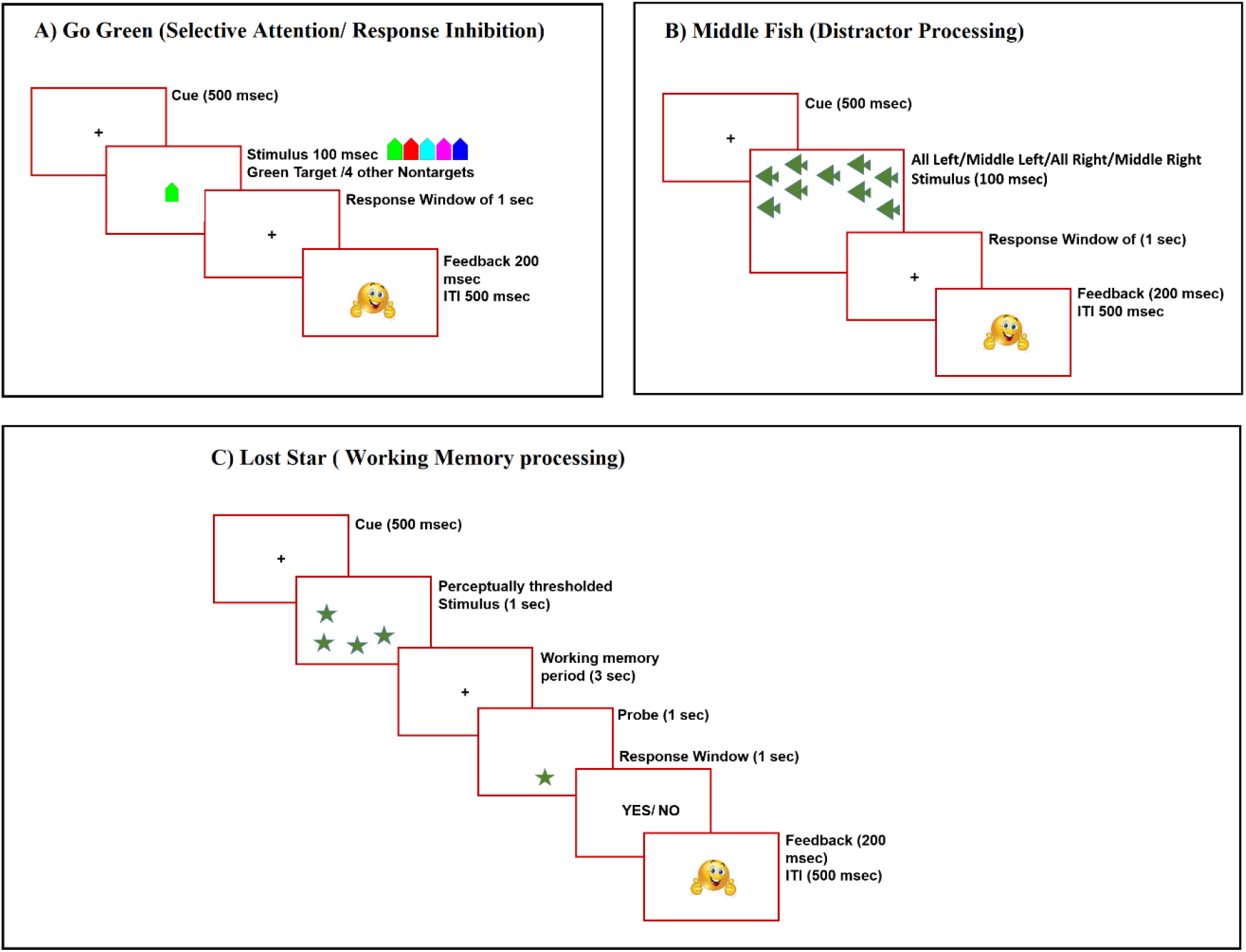
Overview of the various experimental paradigms used to evaluate selective attention, response inhibition, interference processing, and working memory properties. (Balasubramani et al., 2021).

#### 3.1. Experimental Paradigm and Tasks

We modeled three *BrainE* platform tasks. The *Go Green* task was used to study both selective attention and response inhibition. The *Middle Fish* task was used to study processing, and the *Lost Star* was used to assess working memory.

##### Go Green – Selective Attention

Selective attention is tested when the majority of the stimuli are distractors, and very few are targets. In this case, one needs to focus on relevant target stimuli and respond as rapidly as possible. For evaluating selective attention, the *Go Green* (Selective Attention) task was modeled with the test set consisting of 33% green-coloured rocket images and 67% other coloured rocket images. Five different coloured rocket images were trained with appropriate reward processing based on action selection. In this task, when the image of a green-coloured rocket is displayed on the screen, the action ‘GO’ is selected, whereas for all other coloured rocket images, the action ‘NO GO’ is selected. During each test trial, at first, a cue is presented for an interval of 500 ms (milliseconds), followed by the image of the coloured rockets for 100 ms. There is a response window of 1 sec and a feedback window of 200 ms. There is an intertrial interval (ITI) of 500 ms between each trial and the next.

##### Go Green – Response Inhibition

Response inhibition is tested when the majority of the stimuli are targets, and very few are distractors. In this case, one needs to respond to the stimuli most of the time and focus on the distractors and inhibit their response. For evaluating response inhibition, the *Go Green* task was modeled with a test set consisting of 67% green-coloured rocket images and 33% rocket images of other colours. Five different coloured rocket images were trained with appropriate reward processing based on action selection. In this task, when the image of a green-coloured rocket is displayed on the screen, the action ‘GO’ is selected, whereas for all other coloured rocket images, the action ‘NO GO’ is selected. During each test trial, at first, a cue is presented for an interval of 500 ms, followed by the image of the coloured rockets for 100 ms. There is a response window of 1 sec and a feedback window of 200 ms. Between each trial and the next, there is an ITI of 500ms.

##### Middle Fish – DistractorProcessing

For evaluating distractor processing, the *Middle Fish* task was used. In this task, four different images of fish were used to train the model with appropriate rewards. The image with left-facing (right-facing) fish in the center was rewarded with 1 when the action ‘LEFT’ (‘RIGHT’) is selected. Each image consisted of 11 fish, one in the middle and ten others surrounding the middle one. The decision is made based on the middle fish’s orientation (left/right), irrespective of the orientation of the surrounding fish, which act as distractors. During each test trial, a cue is presented for a duration of 500 ms at the start, followed by the array of fish for 100 msec. The images are categorized as all left, middle left, all right, and middle right. All left and all right images have congruent distractors, while middle left and middle right images have incongruent distractors. To win the reward, for all left and middle left fish images, the action ‘LEFT’ should be taken, and for all right and middle right, the action ‘RIGHT’ should be taken. Post-stimulus presentation, a response window of 1 sec, a feedback window of 200 ms and an ITI of 500 ms is programmed between successive trials.

##### Lost Star – Working Memory

For evaluating working memory, the *Lost Star* task was used. In this task, a perceptual image needs to be stored in memory for a brief period, and on presentation of the probe image, action needs to be taken based on whether the location of the star in the probe image matches with that of one of the stars in the perceptual image. One of ‘YES’ or ‘NO’ actions is selected based on the match. A cue is presented for 500 ms at the start of the test, followed by the perceptual image consisting of four stars at random positions on the screen. This test image is presented for one second, followed by a waiting period of three seconds. Then a probe image consisting of a single star is presented for one second. After this, an action must be selected during the response window of 1 sec, followed by a feedback window of 200 ms. Between successive trials, there is an ITI of 500 ms. Eight levels of this task were carried out in (Balasubramani et al., 2021) depending upon the number of stars in the test image. Out of these, in our computational study we have considered only one level (level 0.5 – four stars in the perceptual image) as there was no comprehensive analysis carried out with respect to variation in levels.

### S2. The Architecture

A detailed overview of the model architecture is given in Fig. 2 of the main manuscripts. As already discussed in main manuscript, the model has 5 distinctly identifiable components. 1) Representational System (RS), 2) Memory system (MS), 3) Value Computation System (VC), 4) Explorer Block and the 5) Action Selection System (AS).

#### S2.1 The Representational System (RS)

The input image is presented in the input layer of the RS module (Fig. S2). Convolutional operation is performed on the input image using 16 feature maps, each using 3×3 kernels. The convolutional layers are followed by max-pooling layers that reduce the feature maps by 2×2. After four such stages of convolution and max-pooling, the resultant outputs were mapped to a fully connected layer of size 64×1 which constitutes the output of the RS encoder module. At the decoder end the 1×64 feature output was expanded followed by deconvolutional layers and pooling layers at the end of which the original image was reconstructed back.

One the image is perfectly reconstructed the feature vector output from the fully connected layer of the encoder part of RS is provided as the input the memory system (MS).

**Figure S2:**
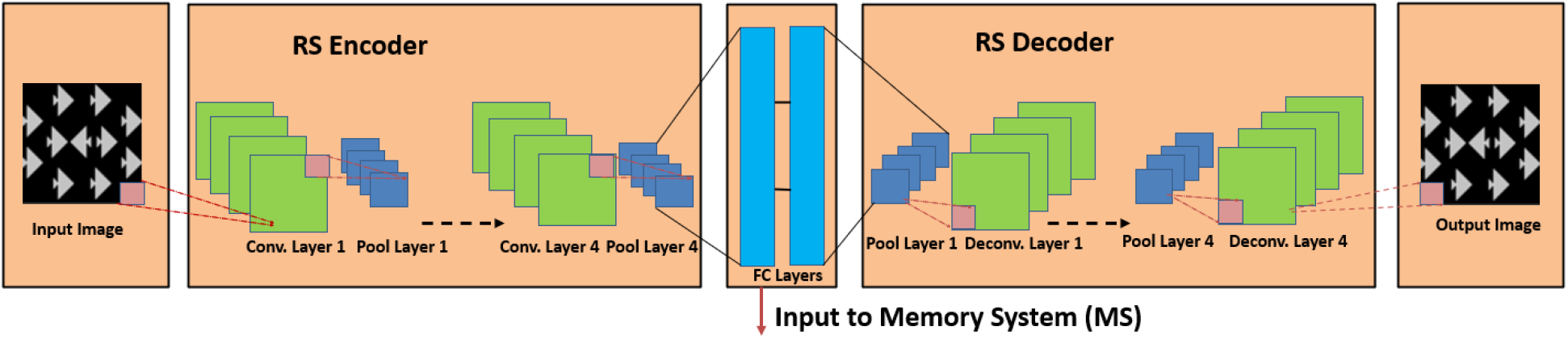
Representational System (RS), with an input and output layer, four convolutional layers and pooling layers and a fully connected layer at the encoder side. The decoder part consists of a fully connected layers, four pooling and deconvolutional layers. When the input image is satisfactorily reconstructed at the output layer, the encoder output taken from the fully connected layer is passed as input to the Memory System (MS).

##### S2.1.1 Flip Flop

The neurons of the memory system (MS) is modelled using J-K Flipflops. There are two inputs to the Flipflop (*J*_*MS*1_, *K*_*MS*1_) and there is one output, *V*^*MS*1^. The inputs for the flip-flop neurons of MS1 and MS2 are given by Equations (1-4). The output of the flip flop neuron can be expressed by Equations (5-6), which is in line with the circuit diagram and the truth table shown in Fig. S3

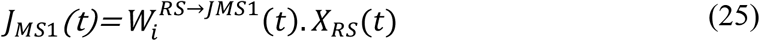

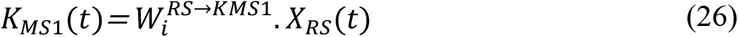

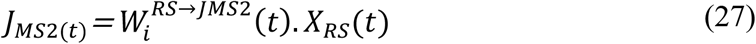

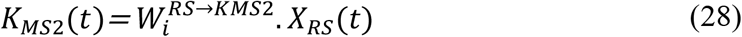

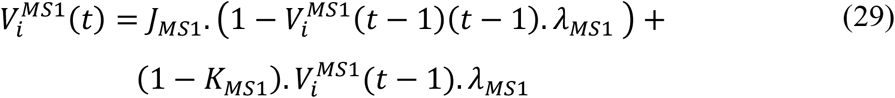

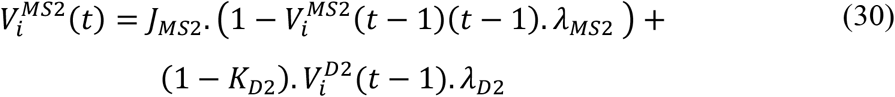

**Figure S3:**
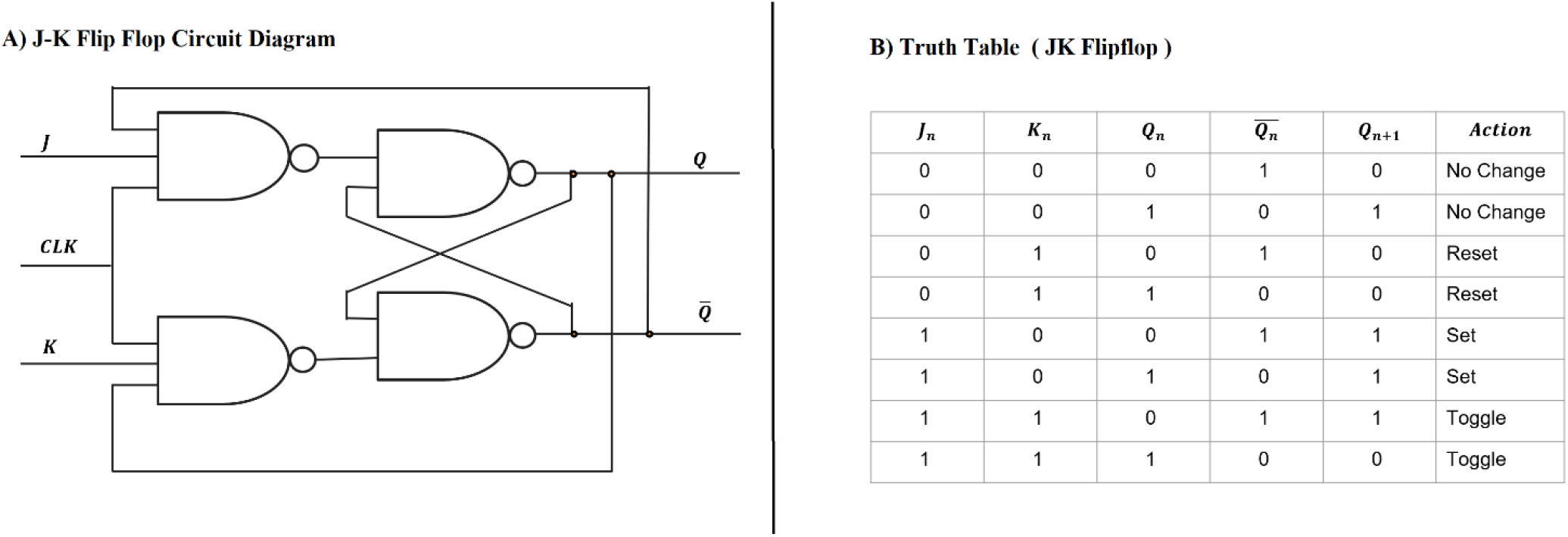
**A) Circuit diagram of JK Flipflop**. J, K inputs to the Flipflop; CLK, Clock input; Q, Output of the Flipflop, **B) TruthTable of the JK Flipflop** If J=K=0, the previous output will be retained, If J=K=1 then the output will be toggled, If (J,K) = (0,1) then the output will be 0, If (J,K) =(1,0) the output will be 1.

### S3. Backward Propagation

The weights between the various modules are updated using the backpropagation algorithm. The learning mechanism in this model is broadly divided into three parts. i) MS1-block to AS weight update, and ii) MS1 block to Value Computation (VC) block weight update, iii) RS to MS1/MS2 block weight update. (Fig. S4).

**Figure S4:**
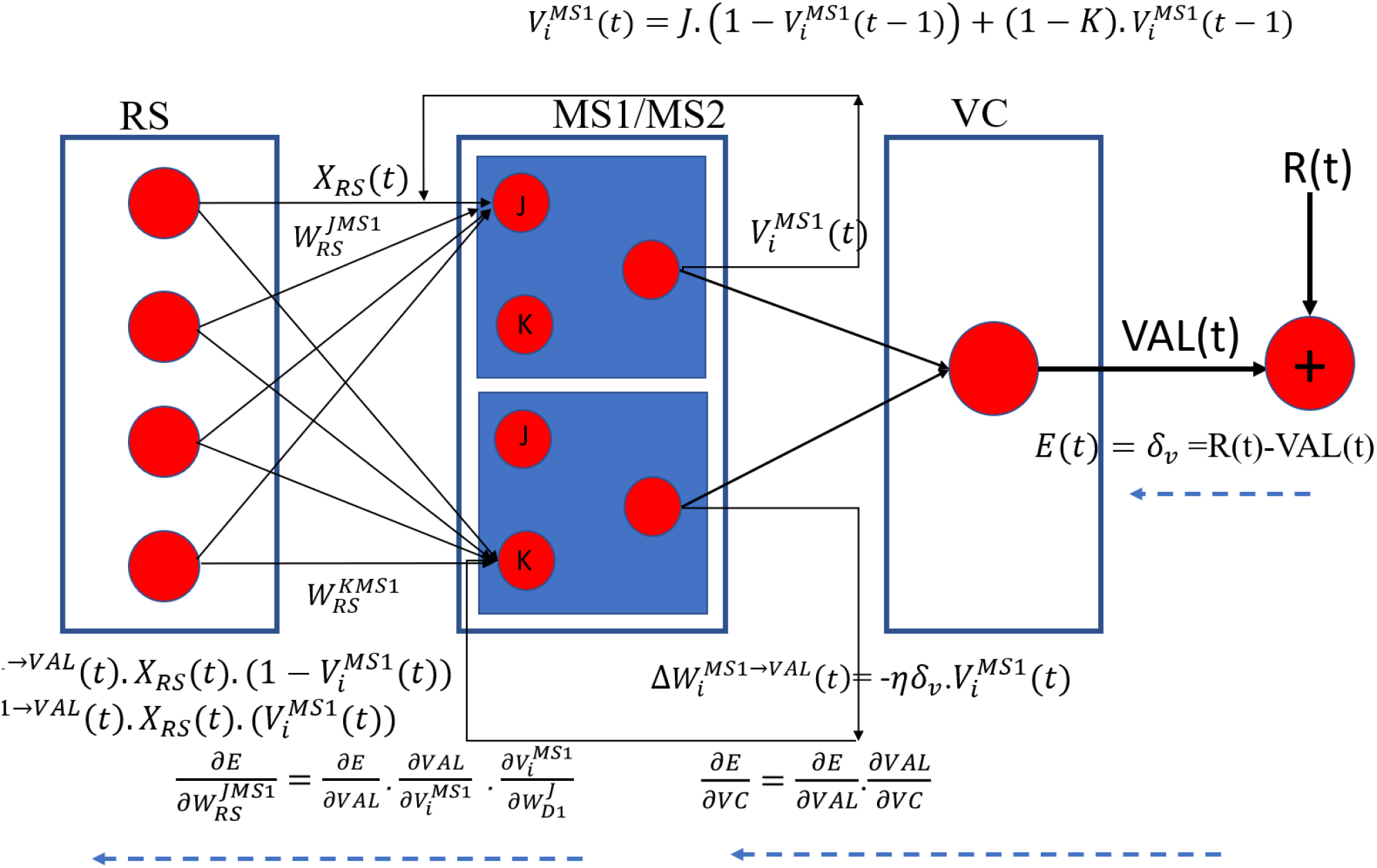
Schematic diagram representing the forward and backpropagation in the GRLDNN model for the circuit Representative System (RS) –> MS1/MS2 block -> VC. VC is the Value computation block; X_RS_, is the input from the representation system (RS) block; MS1/MS2 are the sub-blocks of Memory System (MS)block which forwards the data based on the Modulatory signal from VC.

#### MS1→AS weight update (Q-learning)

The weight update between the MS1 and the AS blocks is governed by Q-learning. The Q table is updated based on the expected reward and the actual reward. The expected reward is high (1) when for a set of given states, the desired action is selected. For example, in case of Go-Green task the Go action is desired when a green rocket is presented, and No-Go action is desired when other coloured rockets are presented. Action Selection block AS is modelled as a race model. Based on the inputs and the D1-AS weights an action is selected out of the possible two. Depending on the action the actual reward will be either 1 or 0. Based on the Expected and the actual rewards the weights are further updated using back propagation using Q-Learning.

The temporal difference error *δ*_*q*_ is used to update the weights between the neurons of the D1 block and the AS block, as shown in Equations (7 to 9). *δ*_*q*_is calculated using the Equation 7. Here r(t) is the reward obtained for selecting the particular action at that time instant, *Q*_*t*_(*s, a*) is the Q value obtained for a particular state action pair and *Q*_*t*+1_(*s, a*) is the future rewards. A discount factor for the future reward given by *γ* is applied to the value that maximizes all possible future actions. The gradient in weight between D1 and AS is found by multiplying this temporal difference error with the output of the D1 block and applying a learning rate *η*_*q*_ as shown in Equation 8.

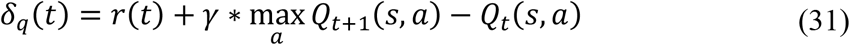

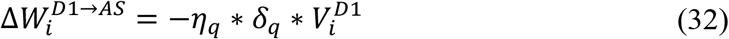

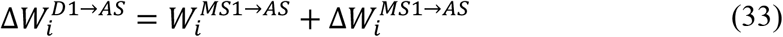

#### MS1 - block → VC weight update

Cortico-striatal weights are updated using 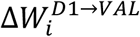, which is the gradient of the error, *E*, with respect to the weights between D1 and M blocks. The Error, *E*, is defined as the difference between the reward obtained for the particular action selected and the Value function as mentioned in Equation. 11. The computation of value function is already shown in Equation. 11 of main manuscript, where the output of the D1 block combines over the connections with respective weights given by *W*^*D*1→*M*^ in order to obtain the Value function (VF).

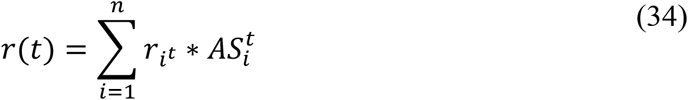

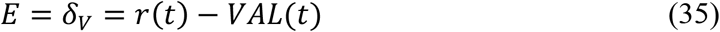

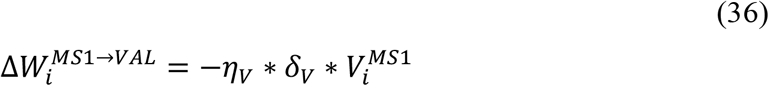

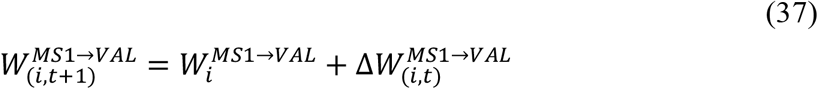

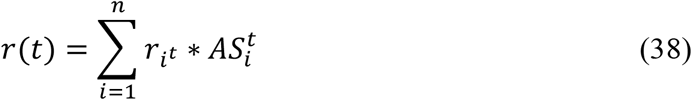

#### RS → MS1/MS2 blocks weight update

The MS1/MS2 sub-blocks that comprise the MS are modelled using the Flip Flop neurons which facilitates the delay and memory properties.

In order to compute the gradients for the weights between the input block and the JK ports of D1/D2 block, let us refer to Fig. 6 The Error, E, in the output that is the mean squared error between the reward received and the value function is propagated backwards through the module M. Here 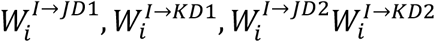 are the weights between the input block,I, and the input J/K ports of the Flip Flops at MS1/MS2 module.

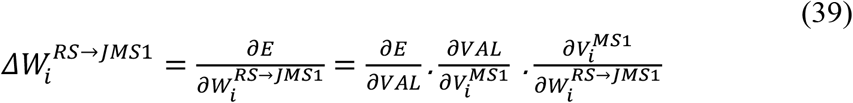

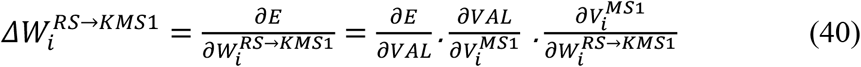

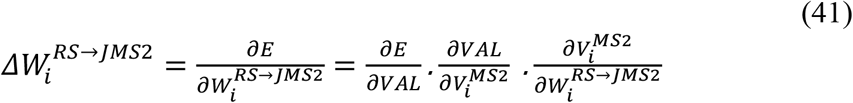

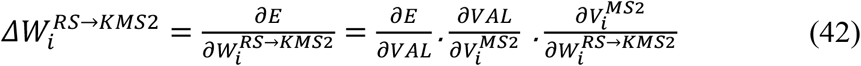

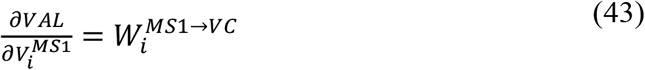

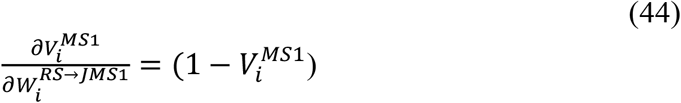

Combining Equations (11,19,20) we can rewrite the Equation (15-18) as Equation (21) and Equation (23-25) respectively.

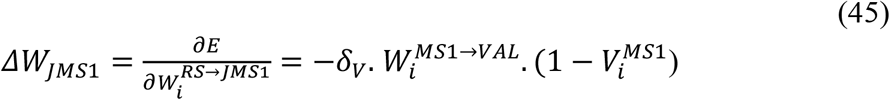

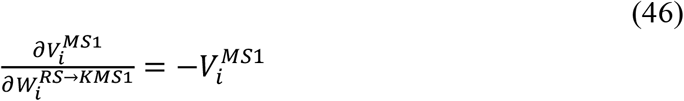

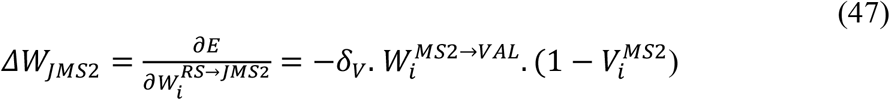

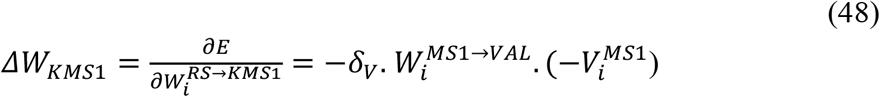

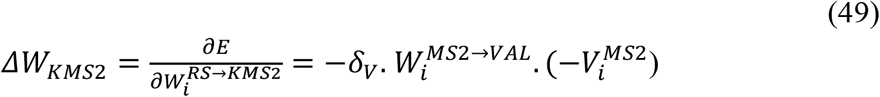

### S3.1 Performance Assessment

The model performance is assessed in terms of the metrics of accuracy, reaction time, speed, consistency, and efficiency, as defined below.

- *Mean Reaction time (RT)* is defined as the number of seconds taken for the winning neuron to cross the threshold.

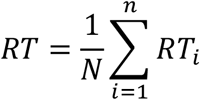
- *Speed* is measured as the logarithm (to the base 10) of the inverse of the reaction time. It indicates the speed of decision-making.

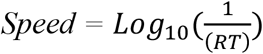
- *Consistency* provides the measure of the consistency of the performance speed.

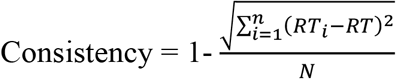
- *Dprime Accuracy, d’ =* z(Hits)-z(False Alarms) (theoretical max 4.65) is the signal detection sensitivity normalized to a 0-1 scale
- *Efficiency* is a composite metric of speed and accuracy; higher efficiency is achieved when there is less speed accuracy trade-off.

The detailed performance results are indicated in the figures of the supplementary section. The below section shows the effect of parameter tuning on performance.

### S4. Results

#### S4.1 Training Phase

During the learning progresses, the value functions keep increasing and approaches the value of 1, which is the maximum value that can be attained. The Q values for the respective states and action pair during corresponding to the correct action is seen to be higher whereas the value corresponding to the wrong action is observed to be lower. Fig. S5 represent the state value functions for various tasks with Fig. S5A representing the Go Green (Selective Attention) task, Fig. S5B representing the Go Green (Response Inhibition) task, Fig. S5C representing the Middle Fish (Distractor Processing) task and Fig. S5D representing the Lost Star (Working Memory) task. Fig. S6 represents the Q value at the end of training for the A) Go Green (Selective Attention), B) Go Green (Response Inhibition), C) Middle Fish (Distractor Processing) and D) Lost Star (Working Memory processing).

**Figure S5:**
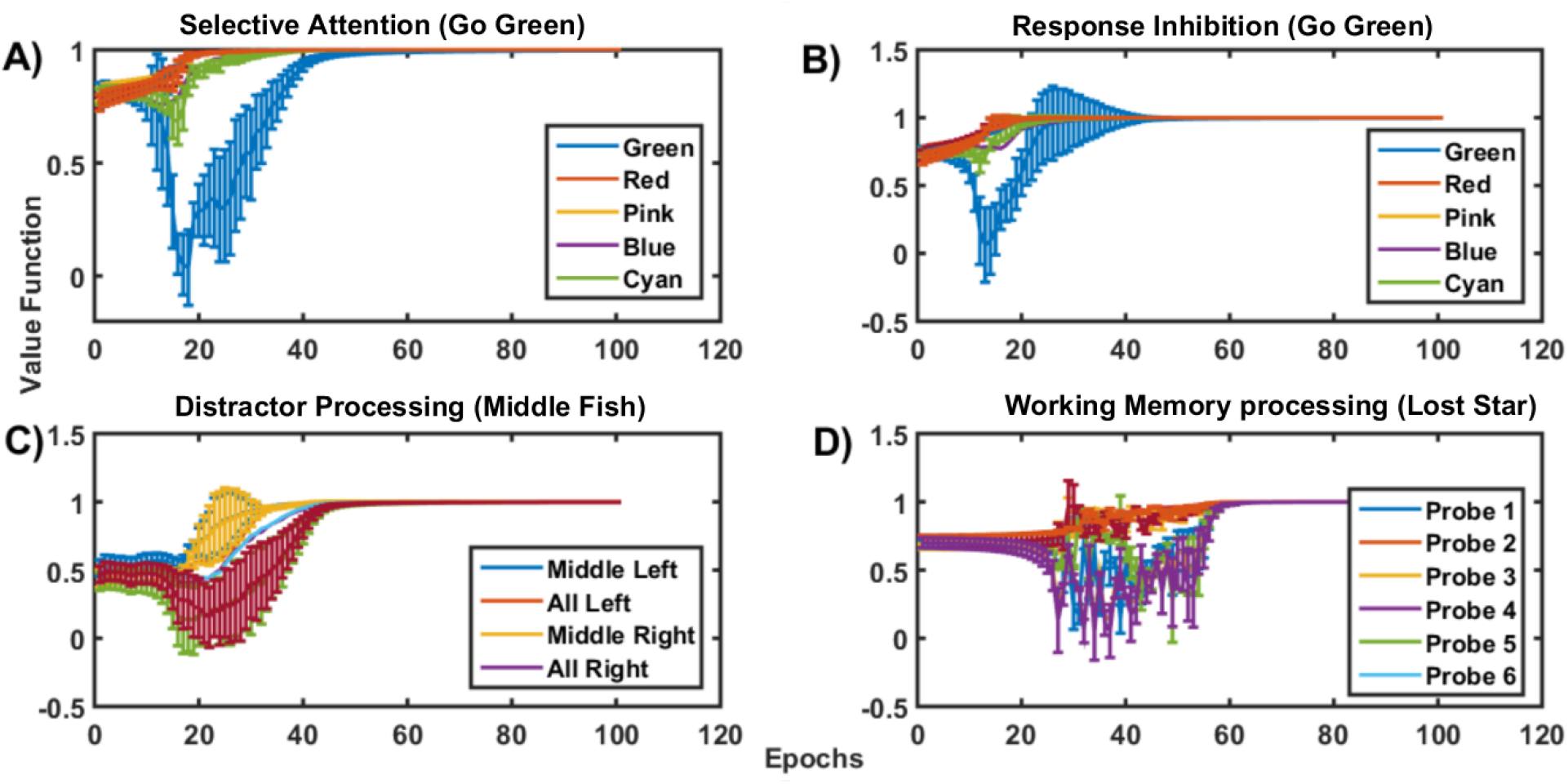
A) The state value functions over the epochs during the training phase for four different tasks as represented in A) The Go Green (Selective Attention) B) The Go Green (Response Inhibition) C) The Distractor Processing task (Middle Fish) and D) The working Memory Task (Lost Star)

**Figure S6:**
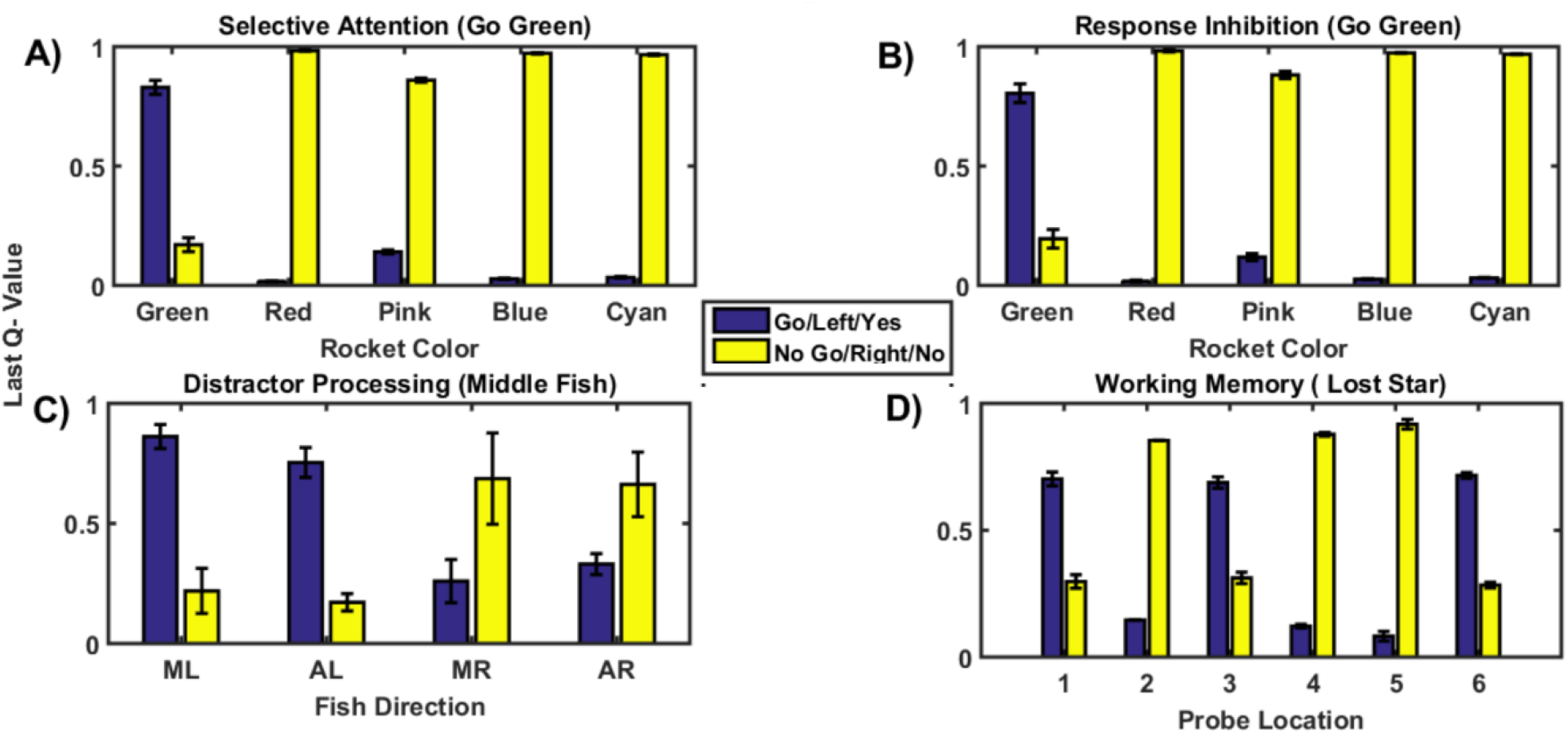
A) The Q values corresponding to the state action pair at the end of the training phase for four different tasks as represented in A) The Go Green (Selective Attention) B) The Go Green (Response Inhibition) C) The Distractor Processing task (Middle Fish) and D) The working Memory Task (Lost Star) is shown. The Q value corresponding to the correct action should be higher and wrong action should be lower. Accordingly, in A and B the Q value corresponding to Go action when green colored rocket is presented is higher and in C the Q value corresponding to the Left action is higher. In case of D, the Q value corresponding to ‘Yes’ action is higher when the star location in the probe image matches that of the perceptual image. The blue bar represents the actions Go/Left/Yes and the yellow bar represents the actions No Go/Right/No.

#### S4.2 Model parameters and Average Performance

We selected the model parameters t *V*_*TH*_ = 0.4 *and ϵ* = 0.05, as the RMSE values were found to be lowest with these parameters when the model performance was compared with that of the average performance of the experimental results. The RMSE values for the four tasks with mdel parameters *V*_*TH*_ = 0.4 *and ϵ* = 0.05, are as given in the Table S1 below.

**Table S1:**
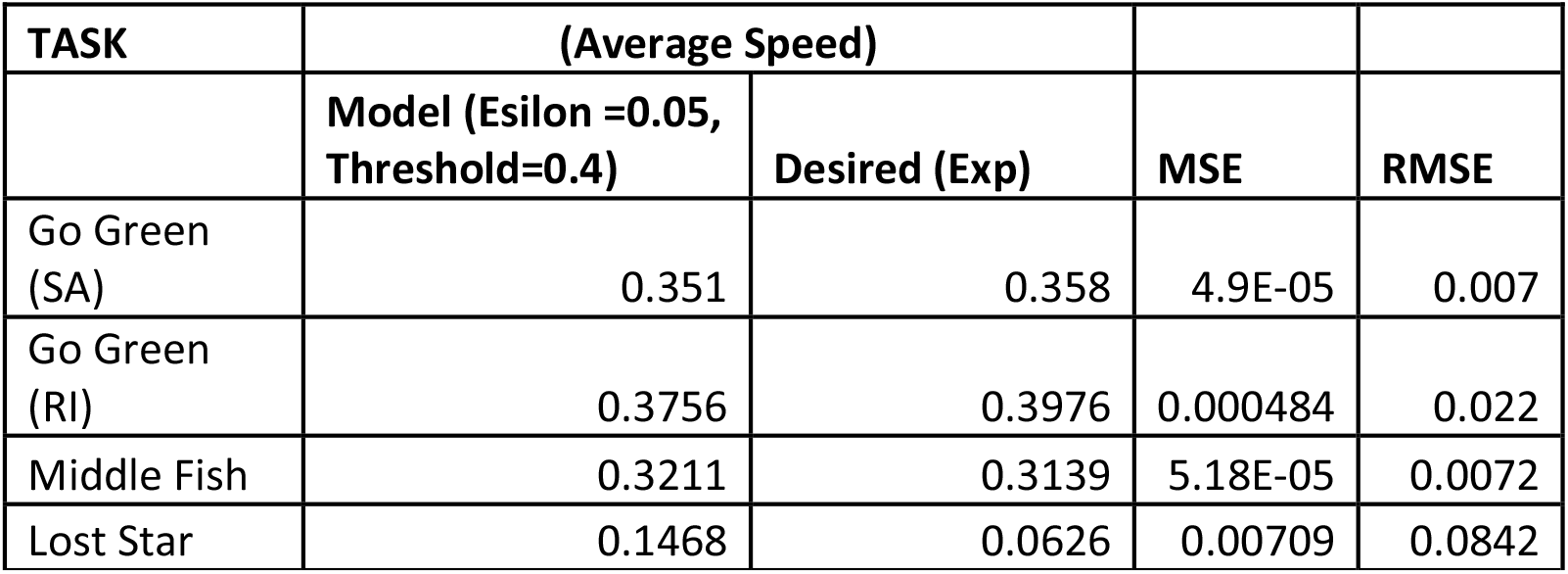
RMSE for different tasks compared to average experimental performance.

#### S4.3 Model parameter Estimation

We modeled the mapping between the experimental parameters (speed, consistency) and the model meta-parameters (*ϵ, V*_*TH*_), tuning using a simple multilayer perceptron model (MLP). Given the input values of speed and consistency, we predict the corresponding values of *ϵ* and *V*_*TH*_ by using a multi-layered perceptron model consisting of one input layer, two hidden layers, and an output layer as shown in Fig. S7. The input layer consisted of 2 neurons whereas each hidden layer consisted of 32 neurons each and after each hidden layer ‘RELU (Rectified Linear Unit)’ activation was used. The output layer consisted of two neurons followed by a sigmoid activation function. The model output gives the predicted values of the meta parameters (*ϵ* and *V*_*TH*_). Hence by varying the values of the two meta-parameters, we would be able to calibrate the GRLDNN model in order to simulate the experimental performance. Twelve data points consisting of different combinations of (*ϵ* and *V*_*TH*_) used to simulate respective speed and consistency values were used for training, and the model was trained for 50000 epochs with the training error in the order of ∼10^−5^ at the end of training. The predicted and desired values of both the threshold and epsilon were closely matched.

**Figure S7:**
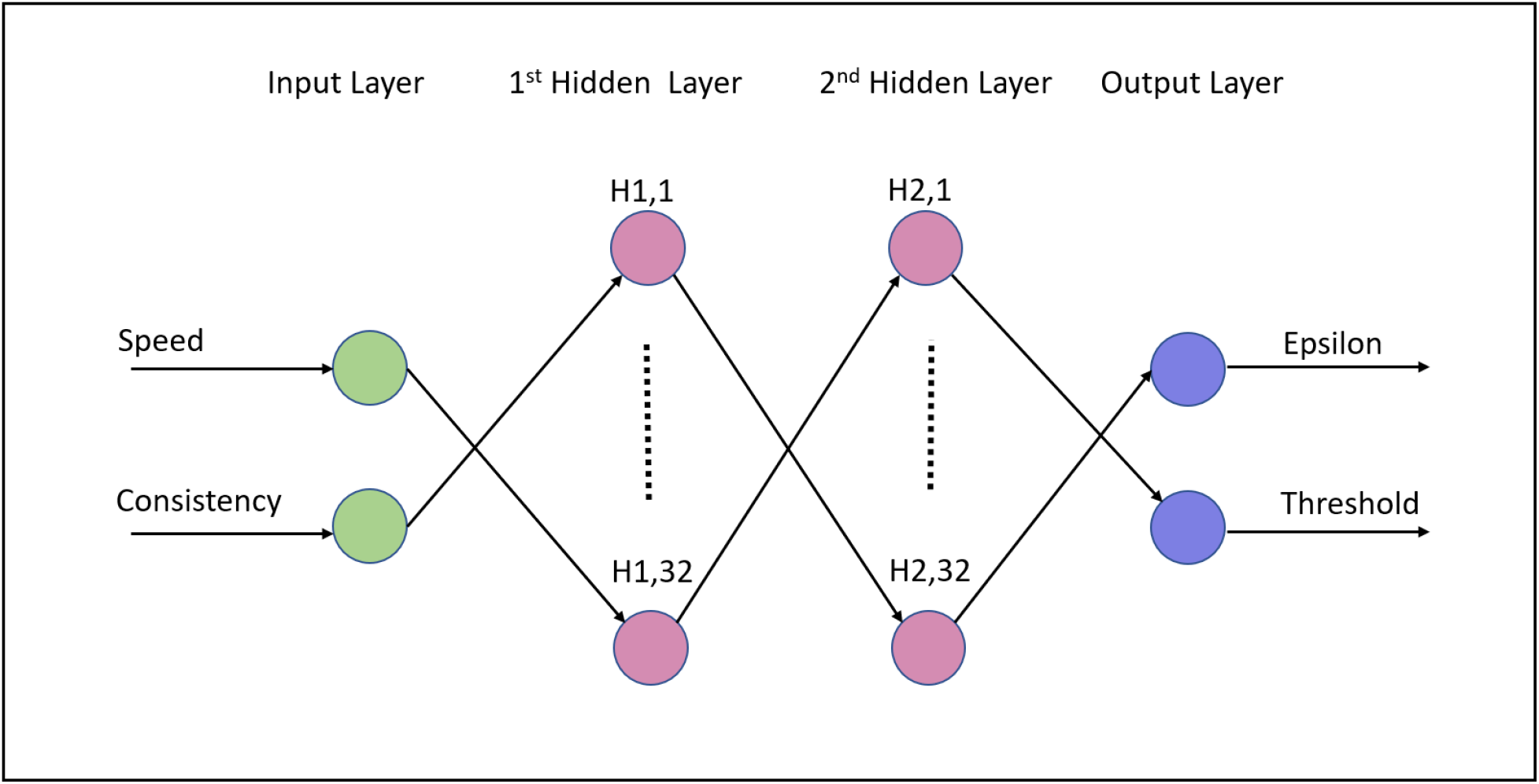
Multi-layered perceptron model to predict the meta parameters with one input layer, two hidden layers and one output layer. Speed and consistency are given as inputs to the model and Epsilon and threshold are predicted. H1,1 to H1,32 represents the neurons in first hidden layer and H2,1 to H2,32 represents the neurons in the second hidden layer.

#### S4.3 Correlation Between Tasks

The empirical performance results of the cognitive tasks indicate strong correlations between selective attention, response inhibition, and distractor processing cognitive functions. This is partly by design, as all of these tasks were set up as speeded stimulus-response tasks. The working memory task showed a very weak correlation with the performance of other tasks. This could very well relate to the differences between the goals of the working memory task, to identify the probe as to whether it matches with a test template but not a particularly speeded response to the probe. Working memory processing also differs from the other core functions of selective attention, response inhibition, and distractor processing in terms of distributed recruitment of brain activation (Smith and Jonides, 1999; RepovŠ and Baddeley, 2006; Konstantinou et al., 2014)

